# The ecological consequences and evolution of retron-mediated suicide as a way to protect *Escherichia coli* from being killed by phage

**DOI:** 10.1101/2021.05.05.442803

**Authors:** Brandon A. Berryhill, Rodrigo Garcia, Joshua A. Manuel, Bruce R. Levin

**Affiliations:** Department of Biology, Emory University, 1510 Clifton Rd NE, Atlanta, GA, 30322, USA; Program in Microbiology and Molecular Genetics (MMG), Graduate Division of Biological and Biomedical Sciences (GDBBS), Laney Graduate School, Emory University, Atlanta, GA, 30322, USA

## Abstract

Retron were described in 1984 as DNA sequences that code for a reverse transcriptase and a unique single-stranded DNA/RNA hybrid called multicopy single-stranded DNA (msDNA). It would not be until 2020 that a function was shown for retrons, when compelling evidence was presented that retrons activate an abortive infection pathway in response to bacteriophage (phage) infection. When infected with the virulent mutant of the phage lambda, λ^VIR^, and to a lesser extent, other phages, a retron designated Ec48 is activated, the *Escherichia coli* bearing this element dies, and the infecting phage is lost. With the aid of a mathematical model, we explore the *a priori* conditions under which retrons will protect bacterial populations from predation by phage and the conditions under which retron-bearing bacteria will evolve in populations without this element. Using *E. coli* with and without Ec48 and l^VIR^, we estimated the parameters of our model and tested the hypotheses generated from our analysis of its properties. Our models and experiments demonstrate that cells expressing a retron-mediated abortive infection system can protect bacterial populations. Our results demonstrate that retron bearing bacteria only have a competitive advantage under a limited set of conditions.

## Introduction

Retrons, DNA sequences that code for a reverse transcriptase and a unique single-stranded DNA/RNA hybrid called multicopy single-stranded DNA (msDNA), were discovered in 1984 (1) and were the first example of a reverse transcriptase coded for by bacteria (2, 3). These elements were initially found in *Myxococcus xanthus*, but subsequently have been observed in a number of bacterial species, including *Escherichia coli* (4, 5). Like CRISPR-Cas, retrons are being employed for genome engineering (5-7), and are capable of doing editing tasks that cannot be done by CRISPR-Cas (8). Also like CRISPR-Cas, the function of retrons was not determined for decades after their discovery and molecular characterization (9). For retrons, the identification of a function came in 2020 when Millman and collaborators presented compelling evidence that retrons mediate an abortive infection (abi) response (10).

With this abi mechanism, the infected cell dies along with the infecting phage (11). From an evolutionary perspective, abi raises interesting questions. Phage-defense systems like envelope resistance, restriction-modification, and CRISPR-Cas are to the advantage of the individual bacteria expressing them. This is not the case for abi; where the defense against phage is through suicide and thereby not to the advantage of the individual bacteria expressing this defense mechanism. To account for the evolution of abortive infection mechanisms, it has been postulated that abi is an altruistic trait which provides an advantage to the collective, clonal population (12, 13). Individual cells that commit suicide due to phage infection prevent these viruses from infecting and killing other, genetically identical, members of their population.

Using agent-based models and experiments with *E. coli* and its phages, Fukuyo and colleagues and Berngruber and colleagues (12, 13), provide a theoretical basis and experimental support for this altruistic suicide hypothesis for the ecological role and evolution of abi. They show that in physically structured habitats, abi can successfully protect populations against invasion by phage, and phage-mediated selection can lead to the evolution of abortive infection.

Here we investigate the conditions under which, by abortive infection, retrons can protect bacterial populations from invasion by lytic phages and the extent to which abi will be selected for in otherwise isogenic bacterial populations lacking this trait. Using a mathematical model of the population and evolutionary dynamics of bacteria and phages in mass culture, we explore the *a priori* conditions under which retron-mediated abi can protect populations of bacteria from invasion by lytic phages, and that phage-mediated selection will enable retrons to become established and maintained in bacterial populations. Using *E. coli* K12 bearing the retron Ec48 described by Millman and colleagues (10) and a virulent mutant of the phage lambda, λ^VIR^, we estimate the parameters of this model and test the validity of the predictions generated from our numerical analysis of its properties for both the protection and evolution hypotheses in liquid and two forms of physically structured habitats.

The results of our experiments support the proposition that in liquid and the two physically structured habitats studied retrons can protect populations of bacteria from infection by phages. Our models and experiments also indicate that the conditions for retrons playing this ecological role are restrictive; the abortive infection mechanism has to be nearly perfect, and there cannot be retron^-^ bacteria that can support the growth of the phages. As anticipated by our models and the theory and experiments reported in (12, 13), in liquid culture in the presence of phage, retrons will not become established in populations of bacteria susceptible to these viruses. Contrary to these earlier studies, in the physically structured habitat of soft agar, abortive infection alone cannot account for the evolution of the retron in populations lacking this element. However, when growing in the structured habitat of colonies on the surface of agar in the presence of phages, a retron^+^ population increases in frequency within a retron^-^ population. Moreover, in all three habitats, in the presence of l^VIR^, envelope resistant mutants emerge in both retron^+^ and retron^-^ *E. coli* populations and ascend to become the dominant population of bacteria.

## Results

### A model of retron-mediated abortive infection

To build the theoretical background to generate hypotheses, design experiments, and interpret their results, we employ a mathematical model. This model is based on the interactions of the populations illustrated in Figure 1, along with the definitions and parameters defined in Supplemental Table 1. In accord with this model, the rates of change in the densities of bacteria and phage and the concentration of the limiting resource are given by the system of time-dependent, coupled differential equations listed in Supplemental Equations (1) – (6).

**Figure 1.**
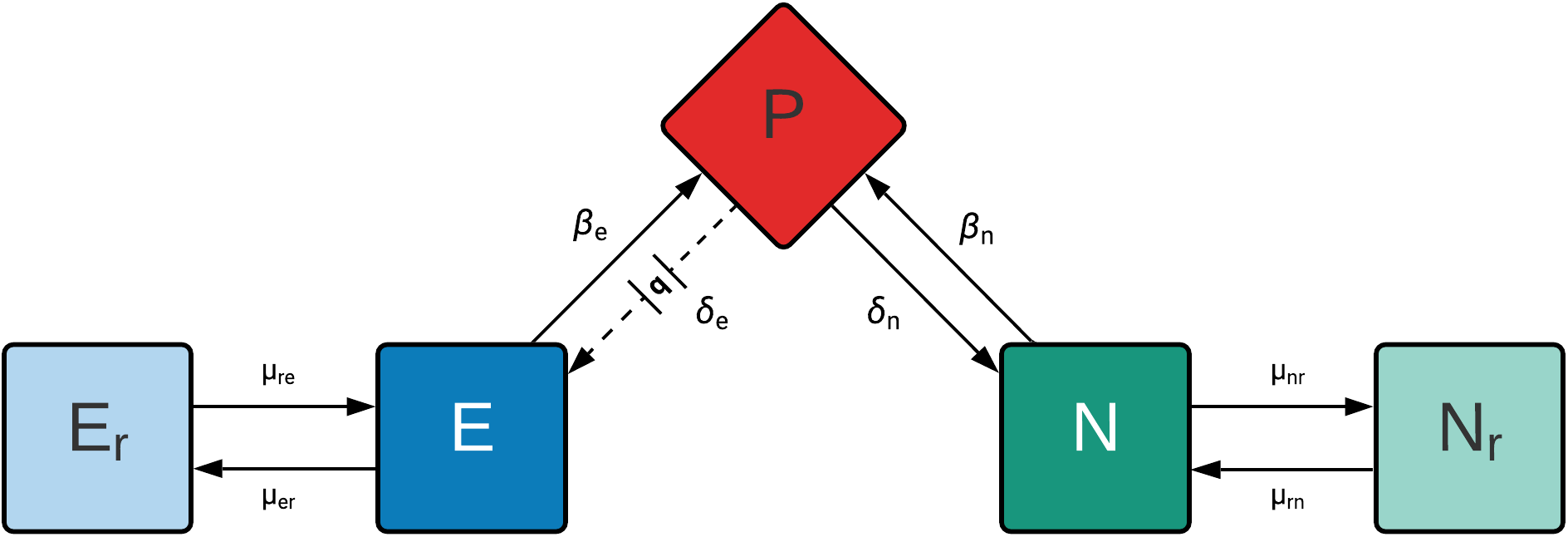
Diagram of a model of the population and evolutionary dynamics of lytic phage and bacteria with and without a retron-mediated abortive infection mechanism. There is a single population of lytic phage, P; a phage-sensitive retron-encoding (retron^+^) population, E; an envelope resistant retron^+^ population, E_r_; a phage-sensitive (retron^-^) population, N; and an envelope resistant retron^-^ population, N_r_. The phage adsorbs to the N and E bacteria with rate constants, *δ*_n_ and *δ*_e_ (ml.cells/hour), respectively. The phage replicates on the N population with each infection producing β_n_ phage particles, the burst size. A fraction q (0 *≤* q *≤*1) of the phages that adsorb to E population are lost and thus do not replicate. The remaining (1 - q) of infections of E produce β_e_ phage particles. At rates μ_nr_ and μ_er_ per cell per hour, the bacteria transition from their respective sensitive to resistant states, and at rates μ_rn_ and μ_re_ they transition from the resistant to their respective phage sensitive states.

### Protection against lytic phages mediated by retrons

We open with an exploration of the population and evolutionary dynamics of retron-mediated abortive infection with an analysis of the ability of retrons to protect populations from predation by lytic phages. For this theoretical and experimental considerations of retron-mediated abi, we present the predicted and observed densities of bacteria and phage when their populations first encounter each other, time 0, and at 24 hours (Figure 2). We performed a theoretical analysis using the parameters and values estimated for this system (Table S1) to determine the minimum efficacy of retron-mediated abi needed to protect populations from phage infection. We tested a range of values of abi effectiveness, q, from 0.95 to 1.0 with steps of 0.001. We found that at least 98% of the infection events have to be aborted by the retron to protect the population from phages (Figure S1). In other words, q must be at least 0.98 for retron-mediated abi to be protective. With these results in mind, we selected q values of 1.00 and 0.95 to illustrate abortive infection success and failure, respectively, Figures 2A and 2B.

**Figure 2.**
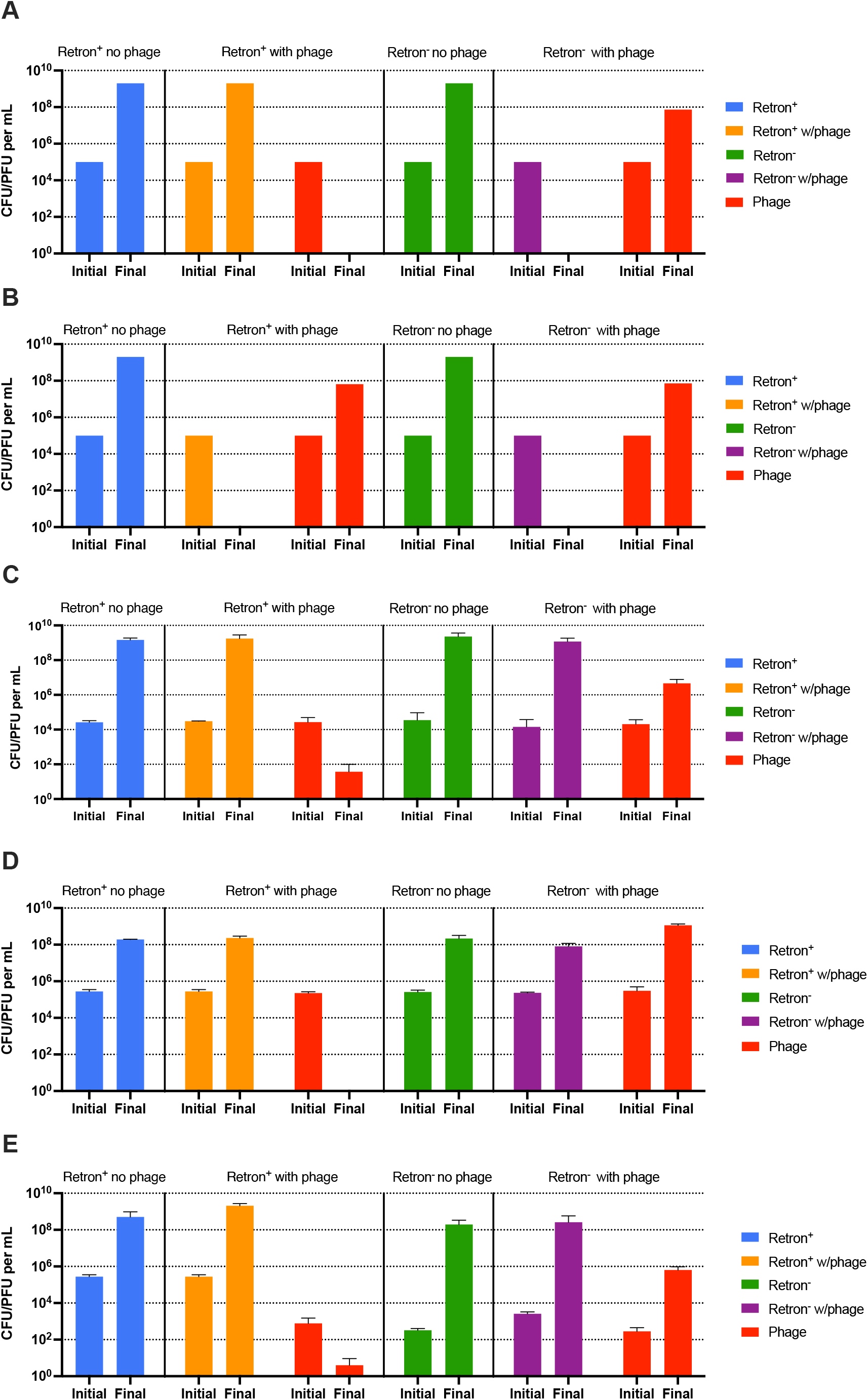
Conditions under which retrons protect bacterial populations from phage infection. Computer simulation results without envelope resistance and experimental results. Shown are the densities of a retron^+^ bacterial population in the absence (blue) and presence (orange) of phage (red) and a retron^-^ bacterial population in the absence (green) and presence (purple) of phage at 0 (Initial) and 24 hours (Final). The parameters used for the simulations where: k=1, e=5×10^−7^ ug/cell, v_e_ = v_n_ = 2.0 h^-1^, *δ*_e_ = *δ*_n_ = 2×10^−7^ h^-1^cell^-1^, β_e_ = β_n_ = 60 phages/cell, μ_nr_=μ_rn_=μ_er_=μ_re_= 0. **A, B-**Computer simulation results with a completely effective (**A**, q=1.00) and incompletely effective (**B**, q=0.95) abortive infection system. **C, D, E-**Protection experiments in liquid (**C**), soft agar (**D**), and with colonies growing on a surface (**E**). Plotted are means and standard deviation of three replicas.

As can be seen in Figure 2A, a completely effective retron-mediated abi defense system (q=1.00) is able to protect a population of retron^+^ bacteria from predation by phages. By 24 hours, the phage population is gone and the retron^+^ population is at its maximum, resource-limited, density. When the retron^-^ populations are confronted with phage, by 24 hours, the bacteria are eliminated and there is a substantial density of free phages. The ability of the retron to prevent the ascent of the phage and protect the bacterial population is critically dependent on the efficacy (q) of retron-mediated abortive infection (Figure 2A, 2B, and S1).

In Figures 2C, 2D, and 2E we present the results of our experimental tests of the retron protection hypothesis presented in Figures 2A and 2B using the retron^+^ *E. coli* Ec48 (10) and a lytic mutant of the phage lambda, λ^VIR^. As a retron^-^ control, we use a λ^VIR^-sensitive *E. coli* C. As anticipated from the model (Figures 2A and 2B), when confronting the retron^+^ population by 24 hours, the λ^VIR^ population is gone or nearly so, and the bacterial density is at the level of a phage-free experiment (Figure 2C, blue bar). The results of the experiment with the λ^VIR^ and the retron^-^ sensitive strain are inconsistent with the prediction of the model. As anticipated by the model, the phage density increased over 24 hours, but contrary to what is expected from the model, the bacteria were not lost, but rather increased to a density similar to that in the absence of phage (Figure 2C). These consistencies and inconsistencies between the experiments and the model obtain regardless of the habitat the experiments were done in. In both soft agar (Figure 2D) and when growing as colonies (Figure 2E) the bacteria were able to survive phage predation regardless of the presence or absence of the retron system, but in populations bearing the retron, the phages were lost.

One possible reason for the survival of the bacteria lacking the retron system in the presence of phages, is that the bacteria recovered at 24 hours from the retron^-^ population are resistant to λ^VIR^. To test this hypothesis, we employed the cross-streak method on colonies isolated at 24 hours to determine their susceptibility to λ^VIR^ (Table S3). By this criterion, the vast majority of the initially sensitive retron^-^ bacteria recovered at 24 hours are resistant to λ^VIR^. This is also the case for the retron^+^ bacteria recovered at 24 hours.

### Invasion of retrons

In Figure 3 we explore the predicted and observed conditions under which a population bearing the retron will be able to increase in frequency in competition with a population of retron^-^ bacteria. Stated another way, we explore the conditions under which the retron^+^ population will be able to invade when rare.

**Figure 3.**
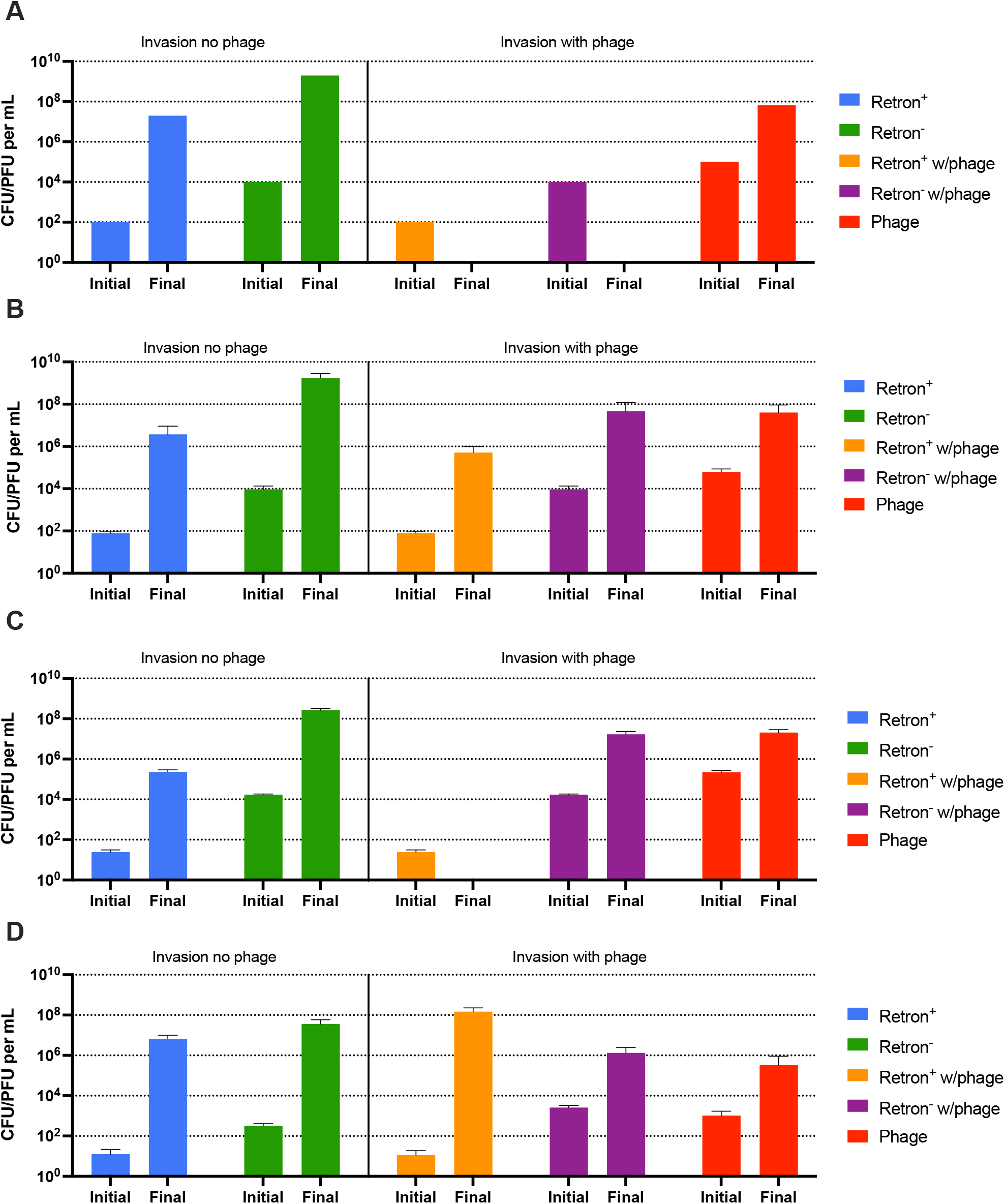
Conditions for the invasion of a retron^+^ population into a community dominated by a retron^-^ population. Computer simulations in the absence of envelope resistance, and experimental results. Bacteria and phage densities at time 0 (Initial) and 24 hours (Final). **Left side:** retron^+^ (blue) and a retron^-^ bacterial population (green) co-cultured in the absence of phage. **Right side:** retron^+^ (orange) and a retron^-^ bacterial population (purple) co-cultured in the presence of phage (red). The parameters used for the simulations where: k=1, e=5×10^−7^ ug/cell, v_e_ = v_n_ = 2.0 h^-1^, *δ*_e_ = *δ*_n_ = 2×10^−7^ h^-1^cell^-1^, β_e_ = β_n_ = 60 phages/cell, μ_nr_=μ_rn_=μ_er_=μ_re_= 0, q=1.00. **A-**Computer simulation results for an invasion condition with a completely effective abortive infection system. **B, C, D-**Invasion experiments in liquid (**B**), soft agar (**C**), and with colonies growing on a surface (**D**). Shown are means and standard deviation of three independent experiments.

As seen in the left side of Figure 3, in the absence of phage in all three habitats, as predicted by the model of mass culture (Figure 3A), the retron^+^ population does not increase in frequency relative to the retron^-^ population, it does not invade when rare. Shown in the right side, and as seen in Figure 2, there is a qualitative difference in the prediction of the model and the observed experiments in the presence of phage (Figure 3B). The model predicts that there will be no bacteria present after 24 hours, yet in all three habitats we find bacteria at the end of the experiment. Consistent with the model (Figure 3A), in liquid (Figure 3B) as well as in soft agar (Figure 3C), the retron^+^ population is not able to invade when rare. Nevertheless, the retron^-^ population at 24 hours reaches a final density similar to the phage-free invasion experiments. In contrast, when growing as colonies in the presence of phage (Figure 3D), the retron^+^ population invades when rare.

To account for the differences in the results of our experiments and those predicted by the model, we performed computer simulations with the model presented in Figure 1, but we now allowed for the generation of phage-resistant retron^+^ (E_r_) and retron^-^ (N_r_) bacterial populations. As noted in Chaudhry *et al*. (14), there is a high rate of generation of λ^VIR^ resistant *E. coli*, suggesting transition rates, μ_er_, μ_nr_, μ_re_, and μ_rn_ of 10^−5^ per cell per hour. With these rates, both the phage and resistant bacteria ascend (Figure 4). The generation of resistance is also consistent with the growth of the retron^+^ and retron^-^ populations in the invasion experiments (Figure 3B). Stated another way, if we allow for phage resistant mutants to be generated in our model, the retron^+^ population can increase in density (Figure 4B, hashed orange), but the dominant population will still be phage-resistant retron^-^ cells (Figure 4B, hashed purple).

**Figure 4.**
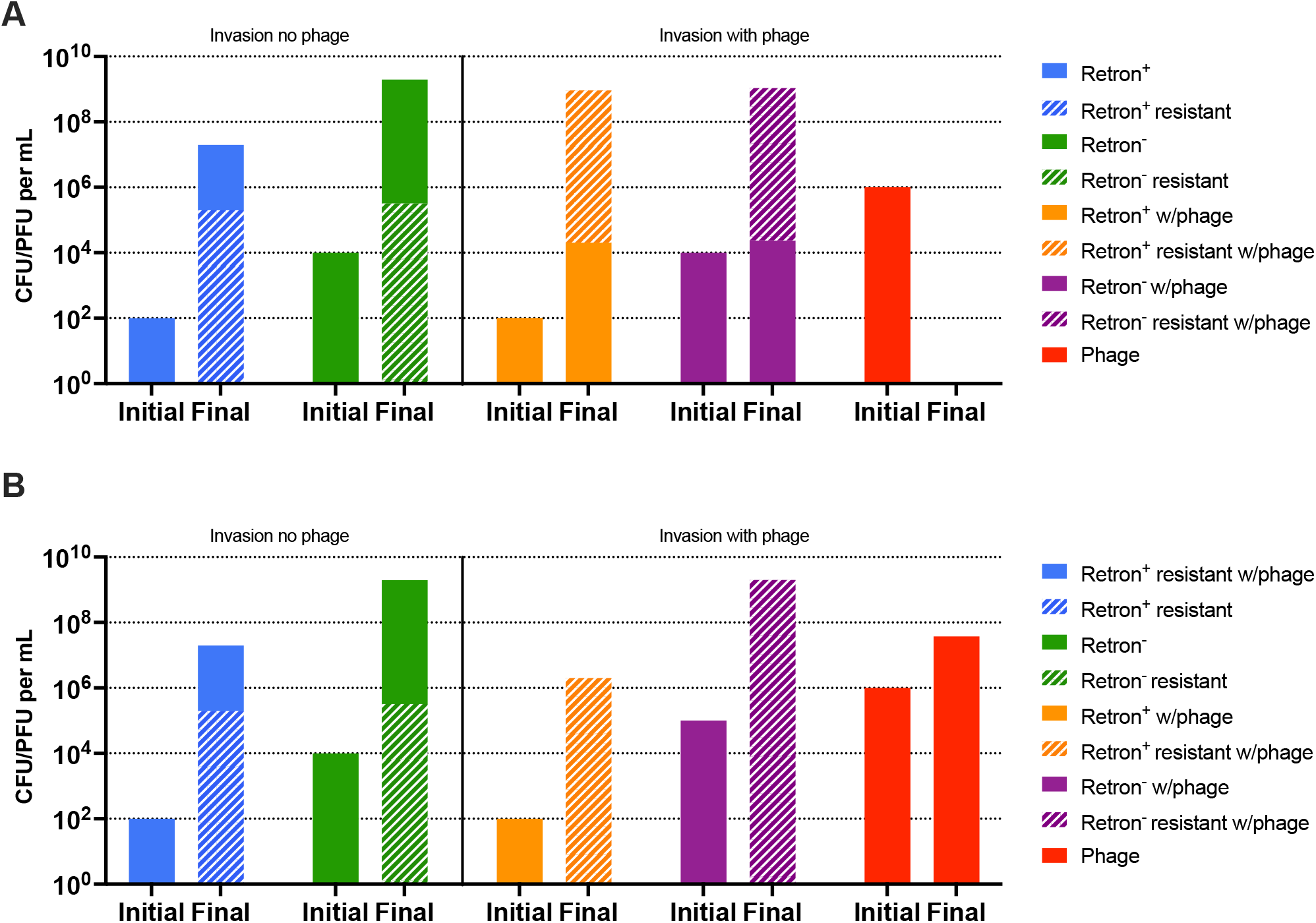
Computer simulations of retron population dynamics with envelope resistance. The simulation conditions are similar to those in Figure 3A but allow for the generation of resistance, μ_er_ = μ_re_ = μ_nr_ = μ_rn_ = 10^−5^ per cell per hour. Densities of bacteria and phage at time 0 (Initial) and 24 hours (Final) for two invasion conditions. **Left side:** retron^+^ (blue) and retron^-^ (green) bacteria in the absence of phage. **Right side:** retron^+^ (orange) and retron^-^ bacteria (purple) in the presence of phage (red). The densities of phage-resistant mutant bacteria are noted by bars with white hashing over the same bar of the sensitive population. The minor population in all cases is overlayed to the greater. **A-**Simulations with a completely effective abortive infection system (q=1.00). **B-**Simulations with a less-than completely effective abortive infection system (q=0.95).

## Discussion

We began the study with two goals: First, to determine the conditions under which retrons will protect populations of bacteria from predation by virulent (lytic) bacteriophages. Second, to determine the conditions under which retron-encoding bacteria will evolve, that is increase in frequency when rare in populations of bacteria without this element. We found that retrons are able to protect populations of retron^+^ bacteria in the presence of phages in mass-liquid culture, structured soft agar, and when growing as colonies on surfaces. Our results also indicate that the conditions under which retron-encoding bacteria are able to invade when rare are narrow. In addressing both of these goals, in our experiments, we found that envelope resistant mutants emerge and become the dominant trait, suggesting that retrons are only one tool bacteria employ as a defense against phage predation.

Our mass action models predict that retron-mediated abi can prevent the populations of bacteria coding for them from being preyed upon by lytic phages, but only if the retron-mediated abi is nearly completely effective, and when there are no other bacterial populations that can support the replication of the phages. The results of our experiments, like those of Millman and colleagues (10) with the retron-encoding abi system in *E. coli* Ec48, and the phage λ^VIR^, are consistent with these predictions. In addition to this protection result occurring in mass, liquid culture, our experiments demonstrate that the protection against lytic phages happens in physically structured populations of bacteria maintained in soft agar or as colonies on surfaces. Our model also predicts if retron-mediated abortive infection is less than 98% effective, with more than 2% of infections being lytic and producing phages, or when there are retron^-^ populations that can support the replication of the phage, retrons will not be able to protect a population from predation by lytic phages. We were unable to test this < 98% efficacy hypothesis experimentally, because our experimental results show that the retron-mediated abortive infection is overshadowed by selection for mutants resistant to the phages. However, since our retron^+^ population was capable of eliminating the phage population when alone, we interpret this to suggest that the efficacy of the Ec48 abortive infection system is over 98%. Even though this retron-mediated abortive infection system is highly effective, when the bacteria are capable of generating envelope resistant mutants, retron^+^ or retron^-^ resistant mutants ascend to dominate the bacterial populations.

Our model predicts in liquid culture that even if a retron-mediated abi defense system is 100% effective in preventing lytic phage replication, and there is an abundance of phages, the retron^+^ population will not be able to evolve by abortive infection alone. Stated another way, when initially rare, the retron^+^ bacteria will not be able to become established in a population of retron^-^ bacteria of similar fitness. Our experiments testing this hypothesis were consistent with this prediction in soft agar as well as liquid culture. As in both habitats, the retron-encoding populations were unable to become established in populations dominated by retron^-^ competitors. However, when growing as colonies in surfaces, we found that the retron-encoding bacteria were able to increase in frequency when initially rare. In all three tested habitats, the *E. coli* population surviving an encounter with λ^VIR^, was dominated by λ^VIR^ resistant mutants.

It had been postulated that in the presences of phages, bacteria with other abortive infection mechanisms can evolve and become established when rare in populations without these mechanisms. Using agent-based models, Fukuyo and colleagues (12) and Berngruber and colleagues (13) predicted that in physically structured communities, where the bacteria are growing as colonies, there are conditions where bacteria with abi systems can invade. Using a constructed abi system, Fukuyo and colleagues (12) found that in the physically structured habitat of soft agar (12, 15), in the presence of phages, their abi-encoding bacteria has an advantage over bacteria without this abi system, but not in a habitat without structure. Similar results were obtained by Berngruber and colleagues (13). In their experiments with *E. coli* growing as colonies in structured environments, depending on the number and size of the colonies, bacteria with their abi system were substantially more fit than the competing population of abi^-^ *E. coli*. In neither of these studies, was the abortive infection system able to evolve in liquid culture. Contrary to these two results, with the retron-mediated abi λ^VIR^ system used in this study, the retron-encoding population was unable to evolve in the physically structured habitat of soft agar, it was only when growing as colonies on a surface that we saw the invasion. At this juncture, we do not know why certain types of physically structured habitats are sufficient to allow for the invasion of the abi trait. Why should soft agar be different than colonies growing on a surface?

## Materials and Methods

### The Mathematical Model

In Figure 1, we illustrate our model of the population dynamics of lytic phage and bacteria with and without a retron-mediated abortive infection system and envelope resistance. There is a single population of phage, P, particles per ml and four bacterial populations of bacteria, E, E_r_, N, and N_r_ cells per ml. The phage sensitive retron population, E, has a functional abi system. Though it also has a function abi system, the E_r_ population is refractory to the phage. The N and N_r_ populations are retron negative, retron^-^, that are, respectively sensitive and resistant to the phage. When a phage infects a bacterium of state E, there is a probability q (0 *≤* q *≤* 1), that the bacteria will die and the infecting phage will be lost. The N population and 1-q of the E population support the replication of the phage while E_r_ and N_r_ are refractory to the phage.

The bacteria grow at maximum rates, v_e_, v_er_, v_n_, and v_nr_, per cell per hour, for E, E_r_, N and N_r_, respectively with the net rate of growth being equal to the product of maximum growth rate, v_max_ and the concentration of a limiting resource, r μg/ml, v_max*_ψ(R) (16), Supplemental Eq (7). The parameter k, the Monod constant, is the concentration of the resource, at which the net growth rate of the bacteria is half its maximum value. By mutation or other processes, the bacteria change states, E→E_r_ and E_r_→E, at rates μ_er_ and μ_re_, per cell per hour, and N→N_r_ and N_r_→N at rates μ_nr_ and μ_rn_.

The limiting resource is consumed at a rate equal to the product of ψ(*R*), a conversion efficiency parameter, e μg/cell (17) and the sum of products of the maximum growth rates of the bacteria and their densities. We assume phage infection is a mass action process that occurs at a rate equal to the product of the density of bacteria and phage and a rate constants of phage infection, δ_e_ and δ_n_ (ml.cells/hour) for infections of E and N, respectively (18). Infections of N by P produce β _n_ phage particles, and the (1-q) of the infections of E by P that do not abort, produce β _e_ phage particles. To account for the decline in physiological state as the bacteria approach stationary phase, R=0, we assume phage infection and mutation rates decline at a rate proportional to Eq.1. The lag before the start of bacterial growth and latent period of phage infection are not considered in this model or the numerical solution employed to analyze its properties.

The changes in the densities of bacteria and phage in this model are expressed as the series of coupled differential equations presented in the supplemental material.

### Numerical solutions – computer simulations

To analyze the properties of this model we use Berkeley Madonna to solve the differential equations (Supplemental Equations (1) – (7)). The growth rate and phage infections parameters used for these simulations are those estimated for *E. col*i and λ^VIR^. Copies of this program are available at www.eclf.net.

### Growth media and strains

Bacterial cultures were grown at 37 °C in MMB broth (LB broth (244620, Difco) supplemented with 0.1 mM MnCl_2_ and 5 mM MgCl_2_). The *E. coli* strain containing the Ec48 retron plasmid was obtained from Rotem Sorek. The sensitive *E. coli* used for controls was *E. coli* C marked with streptomycin resistance, and the Ec48 was marked with ampicillin resistance to differentiate in the invasion experiments. The λ^VIR^ phage lysates were prepared from single plaques at 37 °C in LB medium alongside *E. coli* C. Chloroform was added to the lysates and the lysates were centrifuged to remove any remaining bacterial cells and debris. The λ^VIR^ strain used in these experiments was obtained from Sylvain Moineau.

### Sampling bacterial and phage densities

Bacteria and phage densities were estimated by serial dilution in 0.85% saline followed by plating. The total density of bacteria was estimated on LB hard (1.6%) agar plates. In invasion experiments, diluted samples were placed on LB hard (1.6%) agar plates supplemented with ampicillin (2.5%) or streptomycin (4%) plates to distinguish retron^+^ and retron^-^ *E. coli*. To estimate the densities of free phage, chloroform was added to suspensions before serial dilution. These suspensions were plated at various dilutions on lawns made up of 0.1 mL of overnight LB-grown cultures of *E. coli* C (about 5×10^8^ cells per mL) and 4 mL of LB soft (0.65%) agar on top of hard (1.6%) LB agar plates.

### Resistance Testing with Cross Streaks

Bacteria were tested by streaking in straight lines ten colonies from 24-hour plates across 20 μL of a λ^VIR^ lysate (>10^8^ plaque-forming units [pfu]/mL) on LB hard (1.6%) agar plates. Susceptibility to λ^VIR^ was noted as breaks in the lines of growth. Continuous lines were interpreted as evidence for resistance.

### Growth Rate Estimations

Growth rates were estimated in a Bioscreen C. 48-hour overnights of each strain to be tested were diluted in MMB broth to an initial density of approximately 10^5^ cells per ml. 10 replicas of each strain were loaded into 100-well plates and grown at 37c with shaking for 24 hours taking OD (600nm) measurements every five minutes.

### The Liquid culture experiments

Bacterial overnight cultures grown at 37 °C in MMB Broth were serially diluted in 0.85% saline to approximate initial density and 100 μL were added to flasks containing 10 mL MMB. λ^VIR^ lysate (>10^8^ pfu/mL) was serially diluted to an MOI of ∼1 and 100 μL was added to the appropriate flask. These flasks were sampled for both phage and bacterial initial densities (t = 0 h) and were then grown at 37°C with constant shaking. The flasks were, once again, sampled for phage and bacterial densities (t = 24 h).

### Experiments in soft agar cultures

Bacterial cultures grown at 37°C in MMB and λ^VIR^ lysate were serially diluted in 0.85% saline to appropriate initial densities. The final dilutions were sampled for phage and bacterial initial densities and 100 μL of diluted phage and bacteria were added to 4 mL of LB soft (0.65%) agar and poured into small petri dishes which were grown at 37°C. After 24 hours, the agar was placed into a tube containing 6 mL of saline, vortexed and sonicated in a water bath for 1 hour. These tubes were serially diluted and sampled for final phage and bacterial densities.

### Experiments in SSS

The method employed in the experiments with colonies on surfaces is that developed and employed by Lone Simonsen for a study of the population dynamics of conjugative plasmid transfer in physically structured habitats (19). A volume 4mL of LB agar was pipetted onto a glass microscope slide and allowed to harden. 0.1mL of a phage lysate diluted to the initial densities shown in the figures was placed on one side of the slide and 0.1mL of a bacterial overnight diluted to the initial densities shown was placed on the other side. The liquids were mixed via spreading over the surface of the microscope slide and allowed to grow overnight at 37°C. After 24 hours, the agar was placed into a tube containing 6 mL of saline, vortexed, and sonicated in a water bath for 1 hour. These tubes were serially diluted and sampled for final phage and bacterial densities.

## Acknowledgements

We thank Rotem Sorek and his colleagues at the Weitzmann Institute for providing us with the phage and bacteria used in this study. We thank Kylie Burke, Teresa Gil-Gil, Jake Fontaine, Thomas O’Rourke, Yixuan Peng, Ingrid McCall, Eduardo Rodriguez-Roman, and Andrew Smith for their comments and reviews of this manuscript. We are grateful to Ingrid McCall and David Goldberg, for advice and help with the experimental work. Funds for this research were provided by a grant from the US National Institutes of General Medical Sciences, R35 GM 136407 (BRL).

## Supplemental Material

### Supplemental Equations

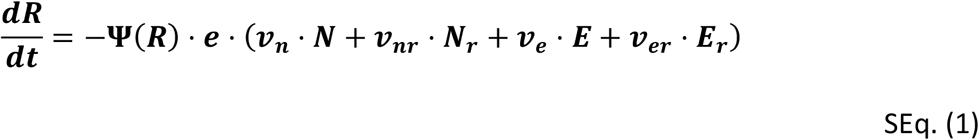

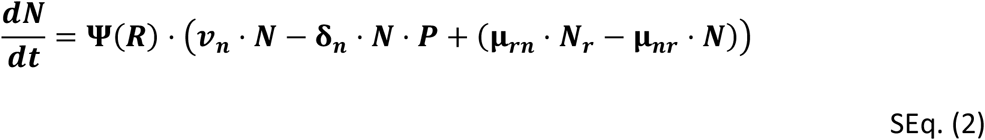

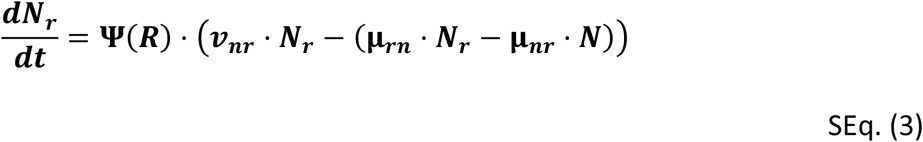

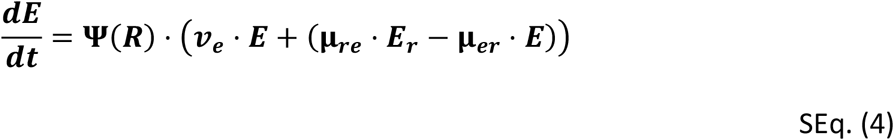

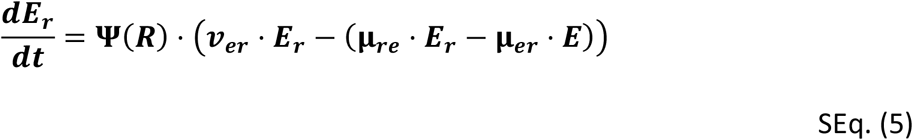

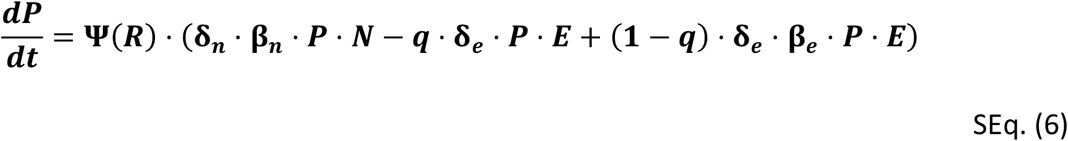

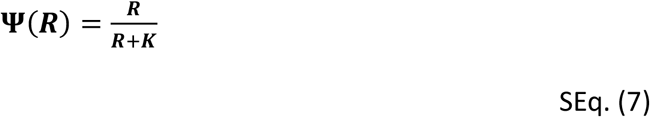

**Figure S1.**
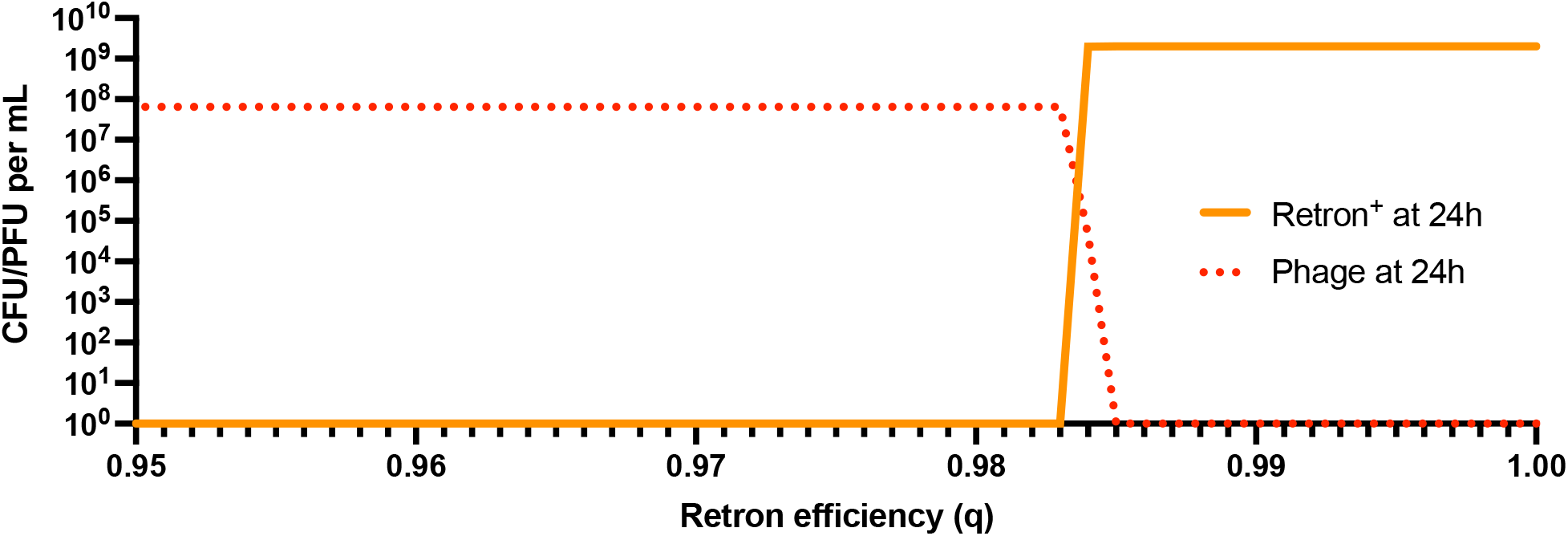
Computer simulation results for the effect of the retron efficiency as values of q. Changes in the densities of a retron^+^ bacterial population in the presence (orange) of phage (red) at 24 hours. Shown are 50 simulations with values of q ranging from 0.95 to 1 with a step size of 0.001 The parameters were the same as in Figure 2.

**Table S1.** Parameter values used in the simulations.

| Parameter | Values (dimensions) | Description | Source (Reference) |
| --- | --- | --- | --- |
| $v_e, v_{er}, v_n, v_{nr}$ | $2.00(h^{-1})$ | Maximum growth rates | This paper (Table S2) |
| $\mu_{nr}, \mu_{rn}$ | $1e^{-5}(h^{-1})$ | Transitions $N \rightarrow N_r, N_r \rightarrow N$ | (1) |
| $\mu_{er}, \mu_{re}$ | $1e^{-5}(h^{-1})$ | Transitions $E \rightarrow E_r, E_r \rightarrow E$ | (1) |
| $\delta_e, \delta_n$ | $2e^{-7}(h^{-1} \cdot mL^{-1})$ | Adsorption rate constants | (1) |
| $\beta_e, \beta_n$ | $60 (PFU \cdot CFU^{-1})$ | Burst sizes | (1) |
| $e$ | $5e^{-7}(\mu g \cdot CFU^{-1})$ | Substrate Conversion Efficiency | (2) |
| $K$ | $1 (\mu g)$ | Monod constant | (2) |
| $q$ | $0 \leq q \leq 1$ | Effectiveness of Abi system | This paper* |

\*Estimated via numerical analysis of the predictions illustrated in Figure S1.

**Table S2.** Growth rate determination.

| Strain (Population) | Symbol | Growth rate ($h^{-1}$)± SE |
| --- | --- | --- |
| *Escherichia coli* Ec48 (E) | $v_e$ | $2.27 \pm 0.06$ |
| *Escherichia coli* Ec48 resistant (Er) | $v_{er}$ | $2.17 \pm 0.02$ |
| *Escherichia coli* C (N) | $v_n$ | $2.37 \pm 0.04$ |
| *Escherichia coli* C resistant (Nr) | $v_{nr}$ | $2.27 \pm 0.02$ |

**Table S3.** Cross-streak results at 24 hours.

| Strain | Plate number | Resistant ratio* |
| --- | --- | --- |
| *Escherichia coli* Ec48 | 1 | 1 |
|  | 2 | 1 |
|  | 3 | 1 |
|  | 4 | 1 |
|  | 5 | 1 |
|  | 6 | 1 |
|  | 7 | 1 |
|  | 8 | 1 |
|  | 9 | 1 |
|  | 10 | 1 |
| *Escherichia coli* MG1655 | 1 | 1 |
|  | 2 | 1 |
|  | 3 | 1 |
|  | 4 | 0.9 |
|  | 5 | 1 |
|  | 6 | 1 |
|  | 7 | 1 |
|  | 8 | 1 |
|  | 9 | 1 |
|  | 10 | 1 |

\* Each colony with a minimum number of 7 from a 10<sup>-7</sup> dilution plate were tested for phage resistance ( $\lambda^{vir}$ ) and the ratio of resistant/total was calculated.

